## Supplementary material for "Host response during unresolved urinary tract infection alters mammary tissue homeostasis through collagen deposition and TIMP1": Sup info

**Affiliations:** [1]Cold Spring Harbor Laboratory; Cold Spring Harbor, NY, USA. [2]Stony Brook University, Graduate Program in Genetics; Stony Brook, NY, USA. [3]CSHL School of Biological Sciences; Cold Spring Harbor, NY, USA.[4]CSHL Undergraduate Research Program; Cold Spring Harbor, NY, USA. [5]Department of Cell Biology and Physiology. School of Medicine in St. Louis. Washington University, St. Louis., MO, USA.[6]SUNY Downstate Health Sciences University, Neural and Behavior Science; Brooklyn, NY, USA. [7]Department of Comparative Medicine, University of Washington; Seattle, WA, USA.[8]Department of Cell Biology. Department of Oncology. School of Medicine. Johns Hopkins University. Baltimore, MD, USA.

### these authors contributed equality

#### Supplemental Experimental Procedures

**Antibodies.** All antibodies were obtained from commercial vendors and used without further purification. *For immunofluorescence staining:* Alexa Fluor 647-conjugated anti-Cytokeratin 5 (EP1601Y, Abcam, cat# ab193895, 0.5 mg/ml, 1:200), Alexa Fluor 488-conjugated anti-β-casein (H-4, SCBT, cat# sc-166530, 0.2 mg/ml, 1:200), FITC-conjugated anti-E. coli antibody (Abcam, cat# ab30522, 5mg/ml, 1:100), anti-human/mouse myeloperoxidase antibody (R&D Systems, cat# AF3667, 0.2 mg/ml, 1:100), Anti-Histone H3 (citrulline R2 + R8 + R17) antibody (Abcam, cat# ab5103, 1 mg/ml, 1:250), Alexa Fluor 568-conjugated anti-goat secondary antibody (Invitrogen, cat# A10037, 2 mg/ml, 1:150) and Alexa Fluor 488-conjugated anti-rabbit secondary antibody (Invitrogen, cat# A21206, 2 mg/ml, 1:150, Picrosirius Red staining kit (Abcam, cat# ab150681), anti-Perilipin 1 polyclonal antibody (Thermo Fisher, cat# PA5-55046, 1:200), Alexa Fluor 488-conjugated anti-Fibronectin (Abcam cat# ab237286, 1:200). *For flow cytometry:* Brilliant Violet 605-conjugated anti-CD45 antibody (30-F11, BioLegend, cat# 103140, 0.2 mg/ml, 3:200), PE/Cy7-conjugated anti-CD11b antibody (M1/70, BioLegend, cat# 101216, 0.2 mg/ml, 1:100) and, Alexa Flour 700-conjugated anti-Ly6G antibody (1A8, BioLegend, cat# 127622, 0.5 mg/ml, 3:200).

**Tissue collection and processing for imaging analyses.** Mammary gland (left inguinal), bladder, kidney, spleen, lungs, liver, intestines, and pancreas were harvested at the experimental endpoint and immediately fixed in 4% paraformaldehyde in 1X PBS (16% solution, Electron Microscopy Sciences, cat# 15711) at 4$^{o}$C overnight, followed by storage in 1X PBS at 4$^{o}$C, until paraffin embedding. Tissue processing, paraffin embedding, sectioning, and mounting onto slides was carried out by the CSHL Shared Histology Core Facility using standard procedures. Mammary gland tissue was sectioned at 7 µm while all other tissues were mounted on to slides in ~5 µm sections. H&E staining were carried out by the CSHL Cancer Center Histology Core Facility using standard protocols.

**Masson's trichrome staining, imaging and quantification.** Masson's trichrome stainings were carried out by the CSHL Cancer Center Histology Core Facility using standard protocols. Stained slides were scanned on an Aperio Light Field Slide Scanner (Leica Biosystems). Image acquisition was carried out with Aperio ImageScope (Leica Biosystems). Image quantification was performed in FIJI (ImageJ) using ten randomly selected images from each gland to quantify the percentage of area stained blue from each image.

**Immunofluorescence (IF) staining.** For the analysis of Cytokeratin 5, β–casein, *E. coli*, Perilipin, and Picrosirius Red, slide-mounted tissue sections were deparaffinized in Xylene (Sigma, cat# 534056) and antigen retrieval was conducted in Trilogy (Cell Marque, cat# 920P-10) by heating for 15 min under high pressure, followed by 5 mins without pressure. Slides were washed in 1x PBS and blocked for 4 hours in a humidified chamber with ~100 µl blocking buffer (10mM Tris-HCl pH 7.4, 100mM MgCl2, 0.5% Tween 20, 10% FBS, 5% goat serum). Tissues were then stained with fluorophore-conjugated primary antibodies in blocking buffer overnight, at 4°C, followed by three alternating washes with 1x PBS (1 min/wash) and blocking buffer (5 min/wash). Nuclear staining with DAPI (Sigma, cat# 10236276001) or propidium iodide (Invitrogen, cat# P3566, 1:1000) was carried out for 10 minutes. Slides were then mounted in ProLong Glass Antifade Mountant (Invitrogen, cat# P36980), cover-slipped and allowed to cure overnight at room temperature. For IF staining of Myeloperoxidase and citrullinated histone H3, a previously published staining protocol was used[1]. After deparaffination and rehydration of the mammary tissue, Tris-EDTA buffer (10 mM Tris Base, 1 mM EDTA, 0.05% Tween 20, pH 9.0) was used for antigen retrieval under low pressure for 8 min. The slides were then washed with cold running water and 1X PBS and subsequently blocked with Fc Receptor Blocker (Innovex Biosciences, cat# NB309) for 30 min followed by 1X Blocking Buffer (0.1% Triton-PBS containing 2.5% BSA and 5% donkey serum) for 1 hour at room temperature. The slides were then washed in 1X PBS and stained with primary antibodies in 0.5X blocking buffer at 4°C, overnight. The slides were then washed and incubated with secondary antibodies and DAPI (10 µg/ml; Invitrogen, cat# D1306) in 0.5X blocking buffer for 1 hour at room temperature. Slides were washed thrice in 1X PBS and water and then mounted and cover-slipped with ProLong™ Diamond Antifade Mountant (Invitrogen, cat# P36961).

**Immunofluorescence (IF) tissue imaging.** IF stained slides were imaged on Zeiss LSM710 confocal microscope with Zen 2012 SP5 software (Zeiss), or on the Leica TCS SP8 confocal microscope (Leica), or on the Zeiss Axio Observer inverted fluorescence microscope using Zen blue 2.0 software (Zeiss) or Leica LAS X software (Leica).

**Immunofluorescence (IF) image analysis.** For Cytokeratin 5 and β-casein analysis, five random field of view were taken per slide that was co-stained with Cytokeratin 5 and β-casein (20x, size 600 x 600, averaging 4, 12 bit, speed 5). In FIJI channels were split, brightness/contrast was altered, but remained consistent across the experiment, and then saved as separate image files. Within these images, percentage of area stained with either Cytokeratin 5 or β-casein across the whole image was quantified in FIJI. For Myeloperoxidase (Mpo) analysis, images were taken under 20x lens with 0.75x zoom-out, and integrated density (MPO to DAPI) were counted across the image after loading images into FIJI.

**Perilipin staining and image analysis**. For perilipin, primary antibody staining was completed overnight at 4°C in the dark and secondary antibody staining was completed for 1 hour at RT in the dark. Images of Perilipin was acquired as described above. For analysis, ten images per stained slide were acquired (40X oil, size 600 x 600, averaging 4, 12 bit, speed 5) and individual adipocytes were manually counted across the image after loading images into FIJI.

**Picrosirius staining, imaging and analysis**. Picrosirius Red staining was conducted using a commercially available kit (Abcam, cat# ab150681) according to the manufacturer's protocol. Briefly, slides were deparaffinized (as described above for IF), rehydrated (by washing twice for 3 min. in absolute ethanol) then stained in Picrosirius Red solution for 1 hour at RT followed by dehydration and clearing (washing twice in xylene) prior to mounting. Then, stained slides were imaged under linearly polarized light on the Zeiss Axio Observer inverted fluorescence microscope using Zen blue 2.0 software (Zeiss). Five images per gland were acquired for calculation of percentage area occupied by thick and thin fibers in FIJI from the acquired images.

**Duct lumen area quantification and count.** Ducts were counted by loading in H&E images of mammary glands into FIJI and counted manually following previously published protocols[2]. Ductal lumen area was quantified after loading H&E images utilizing the tracer and area functions in FIJI.

**Analysis of collagen alignment (Second Harmonic Generation).** Collagen alignment was quantified using CT-FIRE and MATLAB to extract collagen fiber information from two photon second harmonic generation images[3]. One representative field of view was assessed per condition, and each 16X field of view was analyzed by segmenting the image into 25 ROIs of equal size. CT-FIRE was performed on each ROI, and individual fiber information was extracted to determine the degree of alignment per ROI. Relative fiber orientation was determined by finding the difference between each fiber in an ROI and the ROI's mode angle, meaning ROIs with higher alignment would show higher proportions of fibers within 15° to 30° of the mode angle (set to 0°).

**Mammary tissue digestion.** Mammary glands were digested into single cell suspensions using previously published protocols[4–6]. Briefly, inguinal and thoracic mammary glands were harvested, minced and digested for ~90 min at 37°C in RPMI 1640 GlutaMAX (Gibco, cat# 61870127) containing 5% FBS (Corning, cat# 35-010-CV) and 1X Collagenase-Hyaluronidase (Stem Cell Technology, cat# 07912). Digested mammary glands were centrifuged at 2000 rpm for 5 mins and the pellet was cryopreserved in 1 ml of Synth-a-Freeze™ Cryopreservation Medium (Gibco, cat# A1254201), distributed over two cryovials (Corning, cat# 430487), and stored at -80°C for short-term storage and in liquid nitrogen vapor for long-term storage. Viably frozen, digested mammary fragments were thawed in a 37°C water bath, and washed with chilled HBSS (Gibco, cat# 14175103) containing 5% FBS. Single cell suspensions were obtained by incubating the cell pellet with 3 ml of TrypLE Express (Gibco, cat# 12604013) for 3 min followed by an HBSS wash, and subsequent incubation with 1 ml of Dispase (Stem Cell Technology, cat# 07913) containing 40µl DNase I (Sigma, cat# D4263) for 2 minutes. The cell suspension was washed again in HBSS and filtered through a 100 µm cell strainer (BD Falcon, cat# c352360). Centrifugation at 2000 rpm for 5 min at room temperature was used to collect cells after each wash. The cells were resuspended in 1X MACS buffer (1X PBS with 0.5% FBS) and kept on ice prior to staining for flow cytometry.

**Flow cytometry analysis.** Single cells suspended in 1X MACS buffer were stained with appropriate fluorophore-conjugated primary antibodies. Staining was carried out in a 5 ml polystyrene round-bottom tube (Corning, cat# 352054) for 40 min at 4°C in the dark after which the samples were washed in 1X MACS buffer and transferred to a polystyrene round bottom tube fitted with a 35 µm cell strainer cap (Corning, cat# 352235) prior to acquisition on the BD LSRFortessa Dual SORP (BD Biosciences) using BD FACSDiva™ v9 software (BD Biosciences). Data was analyzed using FlowJo™ v10 software (BD Life Sciences). Gating strategy used is described below (Fig. M1).

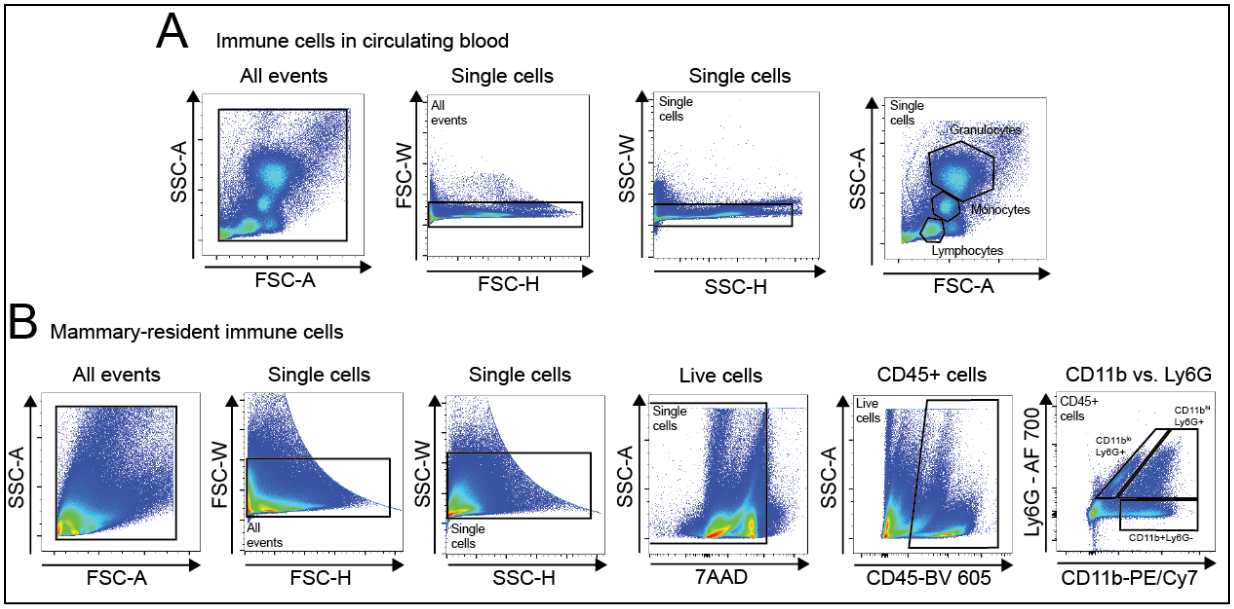

**Fig. M1. Gating strategy for flow cytometry analysis** of **(A)** immune cells in circulating blood and, **(B)** immune cells in the mammary gland.

To assess spectral overlap between the chosen fluorophores, we used single color cell controls and their fluorescence profiles are shown on Fig. M2.

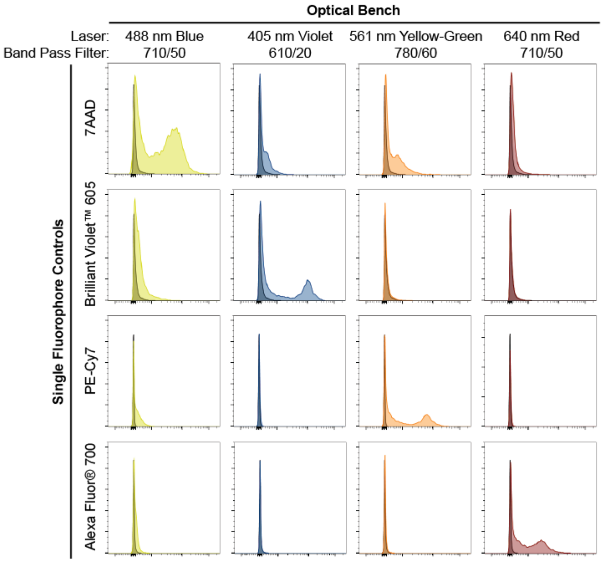

**Fig. M2. Single fluorophore-stained controls for flow cytometry analysis.** Histogram plots for unstained cells (darker histogram) and cell controls stained with individual fluorophore-labeled antibodies (lighter histogram). The plots show fluorescence intensity (x-axis) across all channels and cell count (y-axis). Here, we show that the selected fluorophores had minimal non-specific spill-over and we were able to confidently identify cell populations upon compensation

**Preparation of bacteria for infection.** The uropathogenic *Escherichia coli* (UPEC) strain, UTI89 was obtained as a kind gift from the Hultgren lab at Washington University in St. Louis. UPEC inoculum was prepared as previously described[7]. Briefly, a loopful of UPEC was inoculated from a frozen glycerol stock into 20 ml of LB broth and incubated overnight at 37$^{o}$C, under static conditions. The bacterial culture was then sub-cultured (1:1000) into sterile LB broth and incubated 37$^{o}$C for ~18 hours. The bacteria cultures were centrifuged at 3000 g for 15 min and re-suspended in sterile 1X PBS. Bacteria were enumerated by measuring OD at 600 nm (Ultrospec 7000, GE) and titration on LB agar. The bacterial suspension was diluted in sterile 1X PBS to a concentration of ~2-8 x $10^{9}$ CFU/mL for mouse infection.

**Mouse infections and inclusion criteria.** Mice were infected by delivering active UPEC in 1X PBS (as described above) directly into the murine bladder by transurethral catheterization as described previously[7]. Briefly, catheters were prepared by sheathing a 30 gauge, half-inch needle (BD Precision Glide™, cat# 305106) with polyethylene tubing (PE10, BD Intramedic™, cat# 427401) and were sterilized under UV light for 40 min. The UPEC suspension was loaded into a 1 ml syringe with a tuberculin slip tip (BD, cat# 309659) and fitted with the prepared catheter. Mice were anesthetized by controlled isoflurane inhalation and placed in a supine position; the bladder was voided and the urogenital area was swabbed with 70% ethanol. The catheters were dipped in sterile, non-bacteriostatic lubricant (HR® Lubricating Jelly OneShot®), inserted into the mouse bladder via the urethra and 50 µl (1-4 x $10^{8}$ CFU) of the UPEC suspension was injected into the bladder slowly to minimize vesicoureteral reflux. For controls, mice were administered 50 µl of sterile 1X PBS. For infections in nulliparous mice, 1X PBS (control) or UPEC suspension (UTI) was administered to 7-12 weeks old mice and end-point analyses were conducted at 72 hours or 2 weeks post-infection (p.i.). For infections during lactation (PLI model), mice at lactation day 10 were administered 1X PBS or UPEC suspension. Pups were weaned at 72 hours p.i. to synchronize the initiation of involution. End-point analyses were conducted at involution day 10 (~2 weeks p.i.). Throughout all experiments, mice with bacterial titers above 5 x $10^{4}$ CFU/mL were considered actively infected, and those dropping below this threshold were considered resolved. Animals were tested for CFUs at 48 hours p.i. and endpoint (2 weeks p.i.) and only those actively infected, were analyzed (unless treated with TMS, as described below). At endpoint of 2 weeks p.i., 25/58 nulliparous and 7/15 PLI animals retained infection in our studies.

**Urine collection and bacterial titers.** Mice were restrained by scruffing and the urogenital surface was cleaned by swabbing with 70% ethanol. Free-catch urine samples were collected, by gentle palpation of the bladder, into a clean, sterile petri dish. Urine samples were collected before

UPEC infection, at 48 hours p.i., and at the experimental endpoint. Urine samples were serially diluted, 10-fold, in sterile 1X PBS and spot-plated on LB agar to estimate bacterial counts[7]. Mice with bacterial titers greater than $5 \times 10^4$ CFU/mL were classified as infected.

**Antibiotic treatment.** EQUISUL-SDT antimicrobial oral suspension (Covetrus, cat# 048889) containing Trimethoprim-Sulfamethoxazole (TMS) was added to MediGel® Sucralose cups (2 oz, Clear$H_2O$), at a concentration of 0.48 mg/ml. The TMS cups were placed in the animal cages as the sole source of hydration for the duration of the experiment starting 48 hours p.i.. Urine samples from TMS-treated mice were collected and titrated on LB agar plates to assess UTI resolution.

**Collection of blood cells and plasma.** Blood was collected by terminal cardiac bleeds into EDTA-containing MiniCollect® tubes (Greiner Bio-one, cat# 450475). Plasma was separated from blood cells, by centrifugation at 5000 rpm for 15 min at 4°C, aliquoted and stored at -20°C. The remaining cell pellet was subjected to two rounds of RBC lysis (10 min each) in 1X RBC lysis buffer (BioLegend, cat# 420301) and fixed in 2% PFA for 15-20 min on ice. The fixed cells were washed and resuspended in 1X MACs buffer and stored at 4°C until staining for flow cytometry.

**Cytokine array and ELISAs.** The Proteome Profiler Mouse Cytokine Array Kit, Panel A (R&D Systems #ARY006) was used to profile 40 cytokines and chemokines in plasma and was performed according to manufacturer's instructions. 100 µl of plasma from two mice in each condition were pooled for the assay. Chemiluminescent signal was captured on to an autoradiogram (Lab Scientific, cat# XARALF2025) and developed using a Mini-Med 90, Automatic Film Processor (AFP Manufacturing). The developed films were digitalized by scanning with a Perfection 2450 photo scanner (Epson). The mean pixel intensity of each spot was then measured using FIJI ImageJ[8,9]. ELISAs for detecting corticosterone (Tecan, product# RE52211), Timp1 (Invitrogen, cat# EMTIMP1), and estradiol (DRG International, cat# EIA2693) levels were carried out with commercially available kits according to manufacturer's instructions. Colorimetric readouts were obtained using the SpectraMax i3x Multi-Mode Microplate Detection Platform with SoftMax Pro software (Molecular Devices). Each sample was assayed in duplicate and reported as an average after quantifying the mean density of signal in FIJI. Values are reported relative to the positive control of the kit.

**Bone Marrow isolation, culture, and treatment.** Bone marrow was extracted from the ischium, femur and tibia into HBSS containing 5% FBS by centrifugation at 8000 rpm for 5 min. The cell pellet was resuspended in sterile 1X RBC lysis buffer for 10 mins followed by washing with 1X MACS buffer. The cells were then resuspended in DMEM containing 10% FBS and 1% Pen/Strep and counted on a TC20™ automated cell counter (Bio-Rad). 2 x $10^6$ cells were seeded in each well of a Costar® ultra-low attachment, flat-bottom, polystyrene, 24 well plate (Corning, cat# 3473) in a total volume of 300 µl. *Treatment with plasma:* Bone marrow cells were treated with plasma that had been collected from PBS-treated (uninfected controls) or UPEC-infected mice at a final concentration of 0.5% (v/v). In total, plasma samples from 8 mice/condition were tested in duplicate across three independent experiments. *Sample preparation and flow cytometry:* After the treatment duration, the cells were collected, washed in 1X PBS and fixed with 2% paraformaldehyde for 15-20 min. The fixed cells were washed and resuspended in 1X MACS buffer and stored at 4°C until flow cytometry evaluation. Prior to acquisition, samples were washed in 1X MACS buffer and transferred to a polystyrene round bottom tube fitted with a 35 µm cell strainer cap (Corning, cat# 352235). Flow cytometry sample acquisition was conducted on the BD LSRFortessa Dual SORP (BD Biosciences) using BD FACSDiva™ v9 software (BD Biosciences). Data was analyzed using FlowJo™ v10 software (BD Life Sciences). Gating strategy used is described below (Fig. M3).

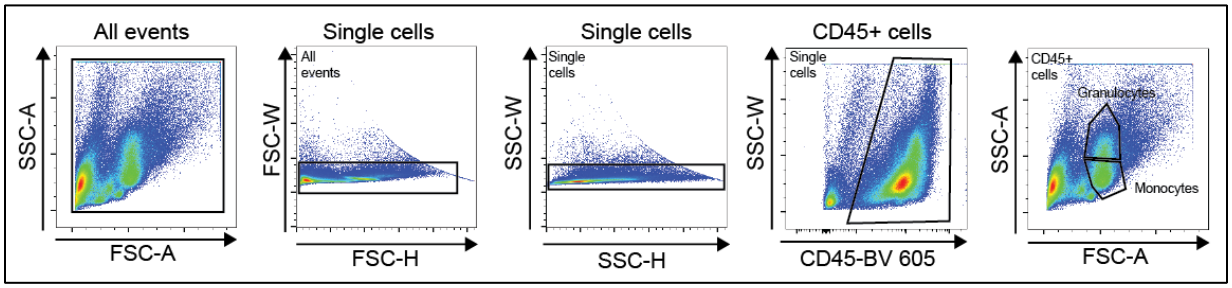

**Fig. M3. Gating strategy for flow cytometry analysis** of bone marrow cells treated with plasma.

**TIMP1 neutralizing antibody treatment.** Mice were infected with UPEC and those with urine bacterial titers exceeding 5 x $10^4$ CFU/mL, at 48 hours p.i., were separated into 2 treatment groups. Either normal goat IgG control antibody (R&D Systems, cat# AB-108-C, 1 mg/ml) or goat anti-mouse TIMP1 neutralizing antibody (R&D Systems, cat# AF980, 0.2 mg/ml) was diluted in sterile, 1X PBS and administered at a concentration of 32 mg/kg, by intraperitoneal injection, every 48 hours up to 2 weeks p.i, the experimental endpoint[10]. In the nulliparous model, of the 37 animals that received UPEC, 27 were confirmed UTI bearing at 48 hours p.i. and of these 27 UTI-bearing animals, 15 were randomly stratified into the IgG control group and 12 into the TIMP1 treatment group. At endpoint, 7/15 IgG animals remained UTI-bearing while 8/12 TIMP1 animals remained UTI-bearing. In the PLI model, of 20 animals that received UPEC, 16 were confirmed UTI-bearing at 48 hours p.i. and of these 16 UTI-bearing animals, 10 were stratified into the TIMP1 treatment group and 6 into the IgG control group. At endpoint, 2/10 TIMP1 treated animals remained UTI-bearing, while 4/6 IgG treated animals remained UTI-bearing.

**CSF-3 neutralizing antibody treatment.** Mice were infected with UPEC and those with urine bacterial titers exceeding 5 x $10^4$ CFU/mL, at 48 hours p.i., were separated into 2 treatment groups. For CSF3 neutralization, mice were administered either rat IgG1 isotype controls (R&D Systems, cat# MAB005) or rat anti-mouse CSF3 neutralization (R&D Systems, cat# MAB414) diluted in sterile PBS at a concentration of 12 μg/mouse in the same course as described above for TIMP1 neutralization[11]. In the nulliparous model, of the 28 animals that received UPEC, 25 were confirmed UTI bearing at 48 hours p.i. and of these animals, 14 were randomly stratified into the CSF3 treatment group and 11 into the IgG control group. At endpoint (2 weeks p.i.), 6/11 IgG animals remained UTI-bearing while 5/14 CSF3 animals remained UTI-bearing.

**scRNA-seq sample preparation.** For all conditions, mammary tissues were collected and digested to a single cell suspension as described above including subjecting the pellet to two rounds of RBC lysis (10 min each) in 1X RBC lysis buffer (BioLegend, cat# 420301). Cell suspensions with a viability greater than 70% were used for scRNA-seq library preparation on the 10X Chromium System (10X Genomics). Single cell libraries were then sequenced (single-end) and indexed using the NextSeq 550 High Output Sequencer (Illumina). Mammary tissue from nulliparous UTI-bearing animas (n=2) yielded a total of 7,100 sequenced cells and post-lactation involution (PLI) control mice (PLI-PBS) (n=2) yielded a total of 11,887 sequenced cells and UTI-bearing (PLI-UTI) (n=2) yielded a total of 9,086 sequenced cells from scRNA-seq. Nulliparous No UTI datasets were previously generated and described[5].

**scRNA-seq data processing and analysis.** scRNA-seq reads were aligned to the mm10 mouse genome assembly using Cell Ranger v.3.1.0 (10X Genomics). Data pre-processing was carried out using Seurat v.4.1.1 using a previously described method[5,12]. Briefly, cells with fewer than 200 or greater than 6000 genes or those with mitochondrial gene content higher than 25% were excluded (Fig. M4).

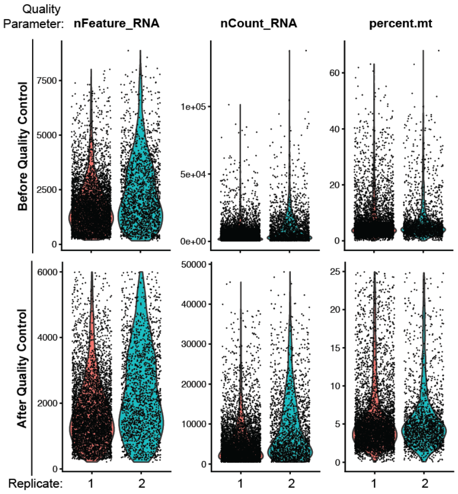

**Fig. M4. Pre-processing of scRNA-seq data for quality control.** Representative violin plots showing nFeature_RNA, nCount_RNA and percent.mt before (top panel) and after (bottom panel) filtering for RNA quality. Briefly, the following quality cut-offs were applied: For nFeatures (left panel), cells outside the range of 200-6000 genes were excluded. For nCount (middle panel), genes represented in lower than 3 cells were removed. For percent.mt, cells with greater than 25% mitochondrial gene content were excluded.

Batch correction was performed on integrated data, using in-built Seurat protocols. Specifically, the FindIntegrationsAnchors() function to identify anchors followed by the IntegrateData() function was used. Principal component analysis (with the top 2000 differentially expressed genes) and unsupervised, hierarchical clustering (resolution = 0.5) was conducted on each Seurat. Gene signature expression scoring was carried out using the AddModuleScore_UCell() function from the UCell package[13]. Statistical analysis on gene expression scores and individual genes was done through stat_compare_means(), a function included in package ggpubr. To quantify cellular cluster population abundance, we utilized Propeller from the Speckle package to quantify statistically significant differences in population abundance between conditions for each cluster[14]. Differential expression between conditions in each cell type was examined through FindMarkers(). This generated gene list was then ranked based on avg_log2FC value and saved as .rnk file for input into Gene Set Enrichment Analysis (GSEA) v4.1 for pathway analysis of Hallmark Terms. Pathways are visualized using ggplot().

**Epithelial cell analysis***.* Cell clusters composed of mammary epithelial cells from nulliparous (NP) or post-lactation involution (PLI) mice, were identified by their expression of genes *Epcam*, *Krt8*, *Krt18*, *Krt5* and/or *Krt14* and extracted from the datasets, re-clustered and re-assessed for quality. Such approach yielded the following cell numbers: NP UTI-bearing = 3,549 cells; NP no UTI = 1,991 cells; PLI control mice (PBS) = 5,350 cells; and PLI UTI-bearing mice = 2,967 cells. These cells were then classified into the three major epithelial cell lineage populations (BM, LASP, LHS) and utilized for further analysis based on previously defined marker genes[15].

**Fibroblast cell analysis**. Cell clusters composed of fibroblasts from nulliparous (NP) or post-lactation involution (PLI) mice were identified by the lack of expression of the genes *Epcam* (epithelial), *Pecam1* (endothelial) and *Ptprc* (immune), and by the expression of the gene *Sparc* as previously described[16,17]. Such approach yielded the following cell numbers: NP UTI-bearing = 1,204 cells; NP no UTI = 84 cells; PLI control mice (PBS) = 2,336 ells; and PLI UTI-bearing mice = 3,607 cells. These cells were then classified into 3 major cellular states according the expression of previously defined maker genes[5].

**CellChat analysis**. Epithelial clusters (BM, LHS, and LASP) and fibroblast clusters (F1, F2, F3) identified above were merged in order to facilitate examination of changes to cellular signaling between epithelial and fibroblast populations using CellChat[18]. In brief, populations were subsetted based on condition (nulliparous: No-UTI, UTI; post-lactation involution: PBS, UTI) and CellChat objects were generated for each condition in each dataset the pre-processed according to previous literature[18]. Visualization of cell signaling networks were generated through functions provided by the CellChat package such as netAnalysis_signalingRole_heatmap() and netVisual_bubble().

**Table M1. Datasets used in scRNA-seq analysis**

| Dataset | Dataset identifier | Reference No. | Figure No. |
|---|---|---|---|
| **Nulliparous (NP)** | | | 2A-I, S3A-F, 4A |
| Condition: No-UTI | SUB8429356 | 5 | |
| Condition: UTI | PRJNA855880 | This study | |
| **Post Lactation Involution (PLI)** | | | 3H-J, S5A-H, S6A |
| Condition: PBS | PRJNA855880 | This Study | |
| Condition: UTI | PRJNA855880 | This study | |

**Table M2. Gene Signatures used in scRNA-seq analysis**

| Gene Signatures | Gene set identifier | Reference | Figure |
|---|---|---|---|

| | | No. | No. |
|---|---|---|---|
| Epithelial lineage identification (scRNAseq) | NA | [5] | 2B, S5B |
| YAP signaling | NA | [19] | 2C-D, 3G |
| ECM signaling | hsa04512 | KEGG | S3A |
| Mechano-sensing | WP4534 | GSEA | S3B |
| Fibroblast Activation Signature | NA | [20] | S5F |
| Neutrophil recruitment | NA | [21] | 4A, S6A |

**Statistical Testing.** Data represents results from at least three independent experiments. All statistical analyses were conducted using GraphPad Prism 7. For comparison of two conditions, an unpaired t-test with Welch's correction was conducted while for comparisons of >2 conditions, one-way ANOVA with Tukey's multiple corrections was used. All collected data was log transformed to adjust for normality. Adjusted p-values lower than 0.05 were considered statistically significant. Across the manuscript $*p \leq 0.05$; $**p \leq 0.01$; $***p \leq 0.001$; $****p \leq 0.0001$. Statistical analysis was supported by the CSHL Biostatistician, Taehoon Ha.

10 Stilley JAW, Birt JA, Nagel SC, Sutovsky M, Sutovsky P, Sharpe-Timms KL. Neutralizing

TIMP1 restores fecundity in a rat model of endometriosis and treating control rats with TIMP1 causes anomalies in ovarian function and embryo development. *Biol Reprod* 2010;**83**:185–94. https://doi.org/10.1095/biolreprod.109.083287.

11 Hollmén M, Karaman S, Schwager S, Lisibach A, Christiansen AJ, Maksimow M, *et al.* G-CSF regulates macrophage phenotype and associates with poor overall survival in human triple-negative breast cancer. *Oncoimmunology* 2016;**5**:1–17. https://doi.org/10.1080/2162402X.2015.1115177.

12 Zheng GXY, Terry JM, Belgrader P, Ryvkin P, Bent ZW, Wilson R, *et al.* Massively parallel digital transcriptional profiling of single cells. *Nat Commun* 2017;**8**:. https://doi.org/10.1038/ncomms14049.

13 Butler A, Hoffman P, Smibert P, Papalexi E, Satija R. Integrating single-cell transcriptomic data across different conditions, technologies, and species. *Nat Biotechnol* 2018;**36**:411–20. https://doi.org/10.1038/nbt.4096.

14 Phipson B, Sim CB, Porrello ER, Hewitt AW, Powell J, Oshlack A. Propeller: Testing for Differences in Cell Type Proportions in Single Cell Data. *Bioinformatics* 2022;**38**:4720–6. https://doi.org/10.1093/bioinformatics/btac582.

15 Stein T, Morris JS, Davies CR, Weber-Hall SJ, Duffy MA, Heath VJ, *et al.* Involution of the mouse mammary gland is associated with an immune cascade and an acute-phase response, involving LBP, CD14 and STAT3. *Breast Cancer Res* 2004;**6**:75–91. https://doi.org/10.1186/bcr753.

16 Bartoschek M, Oskolkov N, Bocci M, Lövrot J, Larsson C, Sommarin M, *et al.* Spatially and functionally distinct subclasses of breast cancer-associated fibroblasts revealed by single cell RNA sequencing. *Nat Commun* 2018;**9**:. https://doi.org/10.1038/s41467-018-07582-3.

17 Kanaya N, Chang G, Wu X, Saeki K, Bernal L, Shim HJ, *et al.* Single-cell RNA-sequencing analysis of estrogen- and endocrine-disrupting chemical-induced reorganization of mouse mammary gland. *Commun Biol* 2019;**2**:1–15. https://doi.org/10.1038/s42003-019-0618-9.

18 Jin S, Guerrero-Juarez CF, Zhang L, Chang I, Ramos R, Kuan CH, *et al.* Inference and analysis of cell-cell communication using CellChat. *Nat Commun* 2021;**12**:1–20. https://doi.org/10.1038/s41467-021-21246-9.

19 Cordenonsi M, Zanconato F, Azzolin L, Forcato M, Rosato A, Frasson C, *et al.* The hippo transducer TAZ confers cancer stem cell-related traits on breast cancer cells. *Cell* 2011;**147**:759–72. https://doi.org/10.1016/j.cell.2011.09.048.

20 Guo Q, Sun D, Barrett AS, Jindal S, Pennock ND, Conklin MW, *et al.* Mammary collagen is under reproductive control with implications for breast cancer. *Matrix Biol* 2022;**105**:104–26. https://doi.org/10.1016/j.matbio.2021.10.006.

21 Neeli I, Khan SN, Radic M. Histone Deimination As a Response to Inflammatory Stimuli in Neutrophils. *J Immunol* 2008;**4**:.
